# Certified Steady-State Parameter-Interval Design for Uncertain Biomolecular Models using a Global Shaving Contractor

**DOI:** 10.64898/2026.05.28.728397

**Authors:** Rudra Prakash, S. Janardhanan, Shaunak Sen

**Affiliations:** Department of Electrical Engineering, Indian Institute of Technology Delhi, New Delhi–110016, India

**Keywords:** Nonlinear system, Parameter design, Interval analysis, Global shaving, Steady-state design, Systems & Synthetic biology

## Abstract

The design of parameter intervals that provably enforce steady-state specifications in biomolecular circuits is challenging due to nonlinear reaction kinetics, parametric uncertainty, and the under-determined nature of steady-state constraints. Most validated approaches either rely on recursive subdivision (set inversion) or may stall due to dependency effects when applied directly in parameter space, limiting scalability in moderate to high dimensions. This paper introduces a global shaving contractor that contracts an initial parameter box by repeatedly applying certified interval-exclusion tests against a prescribed steady-state set. The proposed procedure returns a guaranteed outer enclosure of the feasible parameter set and provides finite-termination guarantees, along with worst-case bounds on the number of inclusion-function evaluations. Case studies spanning low-dimensional motifs and a sixteen-parameter integral-feedback model, including bistability specifications for a CRISPRi toggle switch, demonstrate substantial contraction of design domains without subdivision. The resulting certificates support uncertainty-aware circuit tuning, rigorous parameter screening, and robust design workflows in systems & synthetic biology and related nonlinear dynamical-system design problems.

## 1 Introduction

Robust analysis and design under bounded uncertainty are central themes in control. In synthetic biology, biomolecular circuits implement feedback and decision-making at the cellular scale; however, their dynamical models typically involve nonlinear kinetics and parameters that are only known within broad ranges and may vary across experimental conditions [2, 7]. A recurring design task is therefore to compute parameter sets that are guaranteed to satisfy prescribed steady-state specifications. For example, for bistable architectures such as toggle switches, one seeks parameter ranges that ensure the existence of two distinct equilibria corresponding to the high/low and low/high expression states.

In this work, the steady-state specification is represented by an interval box *X*_*d*_ (or a collection of disjoint boxes in the multi-equilibrium case) together with an initial parameter box *P*_0_. Our objective is to compute a contracted box that is guaranteed to contain all parameter vectors for which the steady-state equations admit at least one solution in *X*_*d*_. Such certificates enable systematic exclusion of infeasible designs and yield reproducible parameter ranges for downstream tuning [21].

A range of tools have been developed to analyze steady states and multistability in biomolecular models. Chemical reaction network theory provides structural conditions for existence and uniqueness of equilibria, as well as robustness properties [9, 25]. Monotonicity and input–output methods certify qualitative behaviors, including multistability and hysteresis, for specific network classes [3, 4, 16]. Methods from real algebraic geometry have been used to characterize parameter-dependent steady-state multiplicity and to discriminate bistable architectures [26]. In addition, sampling-based uncertainty quantification and Bayesian calibration are widely used to identify parameter regimes consistent with data or qualitative specifications [8, 19]. Related estimation and identification problems under uncertainty have also been studied, for example via combined state/parameter estimation with rigorous bounds [5, 6, 14]. However, these approaches are not designed to return guaranteed parameter boxes that certify the existence of a steady state within a prescribed interval *X*_*d*_.

Validated numerics and interval analysis provide a natural framework when explicit guarantees are required. Given a box *Z*, an inclusion function (interval extension) *F* satisfies *f* (*Z*) ⊆ *F* (*Z*) and yields a certified exclusion test: if the zero vector 0 ∉ *F* (*Z*), then *f* (*z*) ≠ 0 for all *z* ∈ *Z* [13, 17, 28]. Throughout, “certified” refers to correctness under standard interval-analysis assumptions, namely inclusion functions evaluated with outward rounding; the corresponding guarantees are stated in Section 2.3.3. This principle underlies interval set inversion based on interval bisection (branch-and-bound) [11, 12] and contractor programming for numerical constraints [13]. In parameter-interval design, generic set inversion typically requires recursive bisection to control over-approximation, which can be prohibitive in moderate and high dimensions [23]. Similarly, interval constraint propagation may stall due to dependency effects, after which subdivision is commonly introduced to recover pruning power [21, 22]. Our previous work [21] addresses the same design objective using subdivision-based interval methods; the present paper instead develops a standalone, certified *no-subdivision* pruning primitive, together with finite-termination and evaluation-complexity bounds and additional case studies.

Within interval constraint satisfaction, *shaving* iteratively tests and removes thin boundary slices of variable domains to enforce local consistency notions (e.g., box consistency) [10, 27]. Traditionally, shaving is used as a local-consistency primitive within branch-and-prune solvers that combine propagation across variables with subdivision. Here we specialize shaving to a design regime in which *X*_*d*_ is fixed *a priori*, the state *x* ∈ *X*_*d*_ is treated as an existential variable, and the objective is to contract only the parameter search box *P*_0_.

We term this specialization *global shaving*: certified contraction of the parameter domain driven by exclusion tests of the form 0 ∉ *F* (*X*_*d*_, *P*_test_) applied to boundary slices *P*_test_ ⊆*P*. The method operates directly in parameter space (without explicit state-space contraction) and admits finite-termination and evaluation-complexity bounds stated in terms of inclusion-function evaluations. Moreover, the pruning primitive composes with simultaneous multi-steady-state requirements via prune-and-intersect, yielding a certified outer enclosure for parameters that satisfy multiple steady-state specifications (e.g., toggle switch design).

The main contributions are as follows:

- We formulate steady-state parameter-interval design with fixed *X*_*d*_ as the computation of a guaranteed outer enclosure of an existential feasibility set in parameter space.
- We introduce a global shaving contractor as a standalone, certified, no-subdivision parameter-space pruning method, including an acceptance rule that commits a shave only when the remaining box is not certified infeasible.
- We establish certification, finite-termination bounds, and worst-case bounds on the number of inclusion-function evaluations, and we evaluate the method on biomolecular case studies ranging from low-dimensional motifs to a sixteen-parameter integral-feedback model.

The remainder of the paper is organized as follows. Section 2 presents the approach, including interval-analysis preliminaries (Section 2.1), the steady-state parameter-design problem (Section 2.2), the global shaving contractor (Section 2.3), and certification/complexity guarantees (Section 2.3.3). Section 3 reports numerical results, Section 4 discusses practical considerations and limitations, and Section 5 concludes the paper.

## 2 Methods

### 2.1 Background

This section summarizes the interval-analysis concepts used throughout; standard references include [13, 17, 18, 28].

An *interval* is a closed, connected subset of the real line, 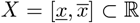 with 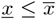. Its diameter (width) is diam 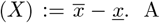. A (hyper-rectangular) *box* is a Cartesian product *Z* = *X*_1_ × … × *X*_*d*_. Let 𝕀ℝ denote the set of closed real intervals; thus *Z* ∈ 𝕀ℝ^*d*^, and we measure box size by diam(*Z*) := max_*i*_ diam(*X*_*i*_).

Interval arithmetic provides rigorous enclosures of the range of elementary operations. For ⋆ ∈ {+, −, ×, ÷} and intervals *X* and *Y* , define the exact range set

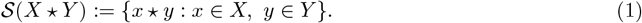

Interval arithmetic computes an enclosure, denoted by *X* ⋆ *Y* , such that 𝒮 (*X* ⋆ *Y*) ⊆*X* ⋆ *Y*. The corresponding endpoint formulas are

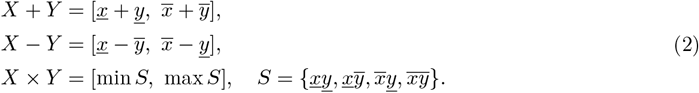

If 0 ∉ *Y* , division is defined by *X ÷ Y* := *X ×* (1*/Y*) with 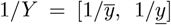. In floating-point implementations, outward rounding preserves the inclusion property [28].

We next recall inclusion functions (interval extensions) and the associated feasibility tests. An inclusion function *F* of *f* satisfies *f* (*Z*) ⊆ *F* (*Z*) for all boxes *Z*. It is *inclusion isotone* if *Z*_1_ ⊆ *Z*_2_ implies *F* (*Z*_1_) ⊆ *F* (*Z*_2_). Consequently, if 0 ∉ *F* (*Z*), then *f* (*z*) ≠ 0 for all *z ∈ Z*, and *Z* can be certified infeasible. If 0 ∈ *F* (*Z*), the test is inconclusive; over-approximation due to dependency and wrapping may prevent pruning even when *Z* contains no solution.

To obtain certified contractions, we employ contractors and domain reduction. Contractor programming provides guaranteed domain-reduction operators that shrink a box while preserving all solutions contained within it [13]. We focus on contractor-only reductions: starting from an initial parameter box *P*_0_, we repeatedly apply certified infeasibility tests to remove boundary slices that fail the inclusion test, thereby contracting the parameter domain.

Although shaving is closely related to box consistency, the objective here is different. In the interval-constraint programming literature, *shaving* enforces a consistency notion (e.g., box consistency) for general numerical constraints by repeatedly testing and trimming boundary slices of variable domains [10, 27]. In contrast, we use shaving in a *design* setting with a fixed steady-state specification *X*_*d*_. Accordingly, shaving is employed to obtain a certified contraction of the *parameter* box by discarding parameter slices (*P*_*test*_) that cannot admit an equilibrium in *X*_*d*_ under the exclusion test 0 ∉ *F* (*X*_*d*_, *P*_test_).

### 2.2 Problem formulation

Consider a parametric ordinary differential equation model

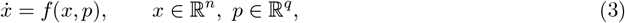

where *f* : ℝ^*n*^ ℝ^*q*^ ℝ^*n*^ is continuously differentiable. Steady states satisfy the algebraic equations *f* (*x, p*) = 0, where 0 ∈ ℝ^*n*^ denotes the zero vector.

The desired steady-state interval box *X*_*d*_ ⊂ ℝ^*n*^ is fixed a priori, and parameter uncertainty is specified by an initial box *P*_0_ ⊂ ℝ^*q*^. The design objective is to compute a contracted parameter box that is guaranteed to contain every *p* ∈ *P*_0_ for which the model admits at least one steady state in *X*_*d*_. A related formulation was studied in our earlier interval-analysis work [21] using a subdivision-based approach. Formally, define the feasible parameter set

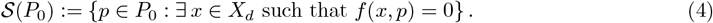

Our goal is to compute a certified outer approximation of 𝒮 (*P*_0_), typically as a contracted box *P*_pruned_ ⊆ *P*_0_ satisfying 𝒮 (*P*_0_) ⊆*P*_pruned_.

To derive mathematically rigorous exclusion tests, we analyze the steady-state equation *f* (*x, p*) = 0 in conjunction with an inclusion-isotone interval extension *F* of *f* (for instance, its natural interval extension) [17, 28]. Since *X*_*d*_ is fixed, the proposed contractor acts directly on parameter boxes. In particular, for any *P* ⊆ *P*_0_ we evaluate the interval image

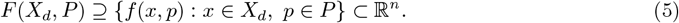

If 0 ∉ *F* (*X*_*d*_, *P*), then no parameter in *P* satisfies the steady-state specification, and *P* can be discarded. More generally, the contractor removes boundary slices of *P* whenever the same exclusion test certifies infeasibility of the slice. Thus, *P*_pruned_ is obtained by domain reduction on *P*_0_ alone, while preserving the guarantee 𝒮 (*P*_0_) ⊆ *P*_pruned_.

In many design tasks, one prescribes *X*_*d*_ and seeks parameter values that satisfy one or more steady-state requirements. When multiple disjoint desired steady-state intervals are specified, it is useful to distinguish an *either–or* requirement (at least one interval contains an equilibrium) from a *simultaneous* requirement (each interval contains an equilibrium, as in multistability design). The following statement captures the simultaneous case used in the positive-feedback example.

#### Theorem 1

(Multiple steady states [21]). *Suppose X*_*d*1_, *X*_*d*2_ ⊂ ℝ^*n*^ *are disjoint and the design requirement is that there exist steady states x*_1_ ∈ *X*_*d*1_ *and x*_2_ ∈*X*_*d*2_ *for the* same *parameter vector p* ∈ *P*_0_. *Let P*_1_ *and P*_2_ *be certified outer enclosures of the parameter sets*

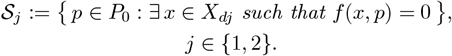

*Then every parameter vector satisfying the simultaneous requirement belongs to P*_1_ *∩ P*_2_.

#### 2.3 Global shaving contractor

We next describe the proposed contractor. The desired steady-state interval box *X*_*d*_ is fixed, and the contractor operates only on a parameter box *P* ⊆ *P*_0_. For notational convenience, define the parameter-to-constraint inclusion map

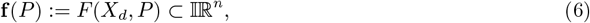

so that **f** (*P*) encloses all steady-state constraint values *f* (*x, p*) for *x ∈ X*_*d*_ and *p ∈ P* .

##### 2.3.1 Inclusion quality: natural interval extension vs. mean-value forms

The power of a certified exclusion test is determined by the tightness of the inclusion function *F*. In the numerical studies, we use the *natural interval extension*, obtained by evaluating the algebraic expressions defining *f* with interval arithmetic and outward rounding. This choice is fully automatic, but it can be conservative when variables are repeated or when nonlinearities induce strong dependency effects.

A common tightening alternative for continuously differentiable constraints is the *mean-value (centered) form* [13, 17, 28]. Let *Z* := *X*_*d*_ *× P* and let *m ∈ Z* denote its midpoint. One may then use the enclosure

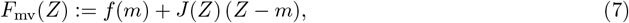

where *J* (*Z*) is an interval extension of the Jacobian of *f* over *Z*. Relative to the natural extension, the mean-value form often reduces over-approximation by linearizing around *m* and bounding the remainder through *J* (*Z*). In global shaving, replacing *F* with *F*_mv_ (or related centered/Taylor-model/affine-arithmetic enclosures) can increase the rate at which boundary slices are certified infeasible, at the cost of additional derivative or bound-propagation computations per evaluation. The case studies below focus on the natural extension to keep the contractor minimal and the cost model *T*_**f**_ (*q*) transparent; tighter enclosures can be incorporated by modularly substituting *F* .

##### 2.3.2 Shaving principle and acceptance rule

Starting from a current parameter box *P*^*\**^, the contractor attempts to remove thin boundary slices. For each coordinate *i* ∈ {1, … , *q*} , it selects a relative slice width Δ = *k* diam 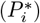 and forms a slice interval *S* at either the lower or upper boundary of 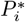. Let *P*_test_ denote the box obtained from *P*^*\**^ by replacing the *i*th coordinate interval by *S*. For vector-valued constraints, we use a componentwise exclusion certificate: if there exists a component *j* such that 0 ∉ **f**_*j*_(*P*_test_), then *P*_test_ contains no feasible parameter and the corresponding slice may be discarded.

When a slice is certified infeasible, we form the remainder box *P*_prop_ obtained by removing *P*_test_ from *P*^*\**^ along coordinate *i*. We commit the contraction only if *P*_prop_ is not certified inconsistent, i.e., if 0 ∈ **f**_*j*_(*P*_prop_) holds for every component *j*. Within each directional shaving loop (lower or upper face), the contractor may remove multiple consecutive slices: whenever a shave is committed, the interval 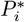 is updated, the slice width Δ is recomputed, and shaving continues until the exclusion/acceptance tests fail or the slice width drops below the tolerance.

The overall procedure terminates when a full sweep over all coordinates yields no contraction, or when all candidate slice widths satisfy *k* diam 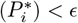.

Algorithm 1 summarizes the resulting global shaving contractor.

##### 2.3.3 Certification and complexity

###### Theorem 2

(Certification). *Assume F is an inclusion function of f* (*x, p*), *that is*, {*f* (*x, p*) : *x ∈ X*_*d*_, *p ∈ P*} ⊆ *F* (*X*_*d*_, *P*) *for every box P* ⊆ *P*_0_. *Let P be the input parameter box to Algorithm 1 and let P*^*\**^ *be its output. Then P*^*\**^ ⊆ *P and every feasible parameter p* ∈ *P (that is, a parameter for which there exists x* ∈ *X*_*d*_ *satisfying f* (*x, p*) = 0*) is also contained in P*^*\**^. *Moreover, if Algorithm 1 returns* ∅, *then no parameter in P satisfies the steady-state specification*.

###### Algorithm 1

Global shaving contractor

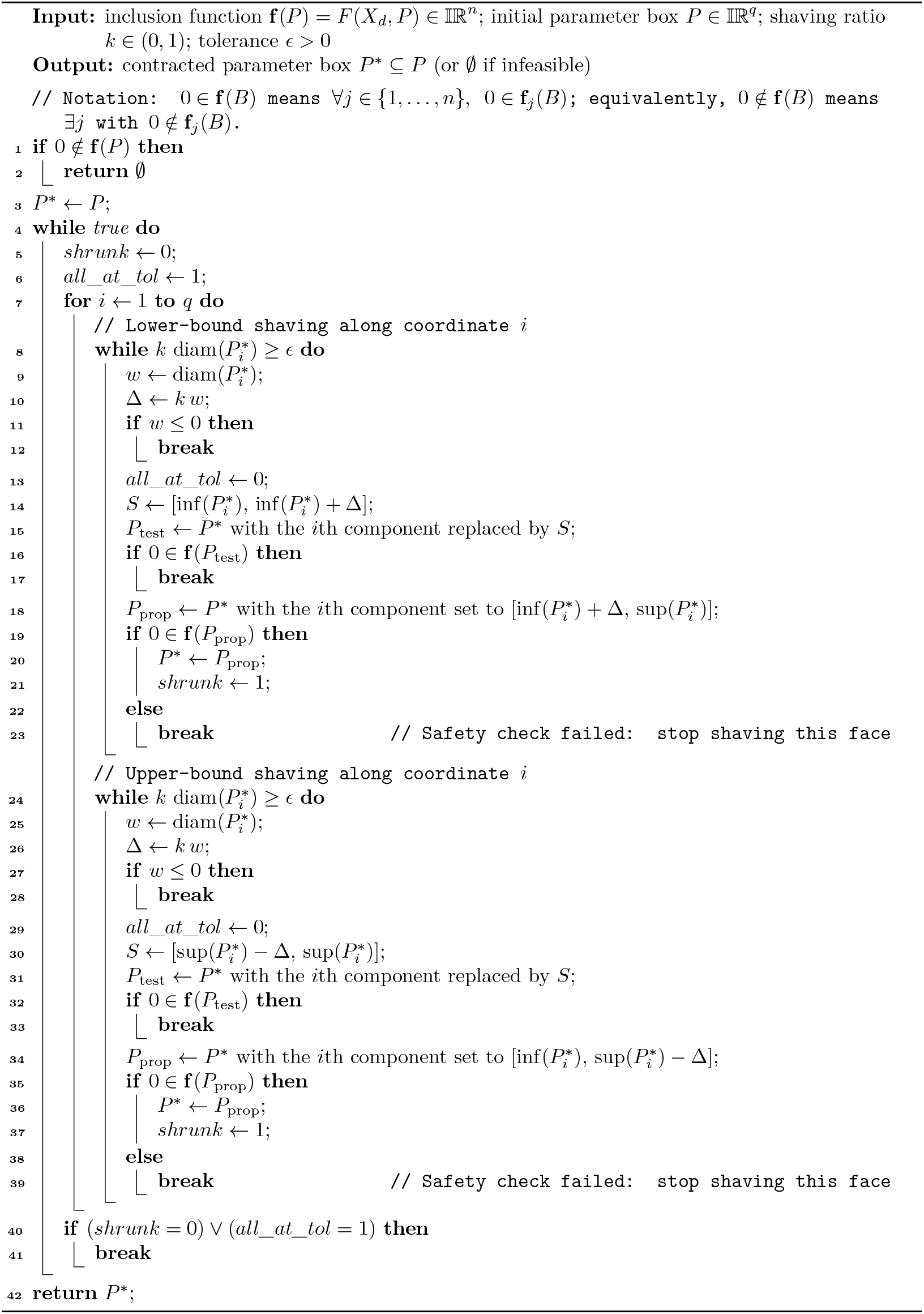

*Proof*. If 0 ∉ **f** (*P*) = *F* (*X*_*d*_, *P*), then by the inclusion property there exists no *x* ∈ *X*_*d*_ and no *p* ∈ *P* such that *f* (*x, p*) = 0, and thus *P* contains no feasible parameter. Hence returning ∅ is correct.

Otherwise, consider any shaving step in which a slice box *P*_test_ is removed. By construction, removal occurs only when 0 ∉ **f** (*P*_test_) = *F* (*X*_*d*_, *P*_test_). Again by the inclusion property, this implies that *f* (*x, p*) ≠ 0 for all *x* ∈ *X*_*d*_ and all *p* ∈ *P*_test_, so *P*_test_ contains no feasible parameter and can be discarded. The additional acceptance test 0 ∈ **f** (*P*_prop_) can only prevent a contraction and therefore cannot remove feasible parameters. Repeating the argument over all successful shaving steps proves that no feasible parameter in *P* is removed, and thus 𝒮 (*P*) ⊆ *P*^*\**^.

□

###### Theorem 3

(Finite termination bound). *Fix a coordinate i and consider one directional shaving loop (lower or upper) in Algorithm 1. Assume the corresponding shaving test succeeds at every iteration until the slice width becomes smaller than ϵ. Then the number of successful shaves on that side is at most*

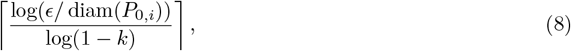

*where P*_0,*i*_ *denotes the ith interval of the initial parameter box P*.

*Proof*. Each successful shave reduces the width of the active interval by a factor (1 *− k*). The bound follows by solving (1 *− k*)^*t*^ diam(*P*_0,*i*_) *< ϵ* for *t*.

□

###### Theorem 4

(Worst-case time complexity). *Consider Algorithm 1 applied to a q-dimensional parameter box P* = *P*_0,1_ ×· · · × *P*_0,*q*_. *Assume that one evaluation of the interval inclusion function* **f** (·) *over a q-dimensional box has cost T*_**f**_ (*q*). *Define*

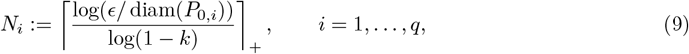

*where* ⌈·⌉_+_ *denotes* max{0, ⌈*·*⌉}. *Then, in the worst case, Algorithm 1 performs at most*

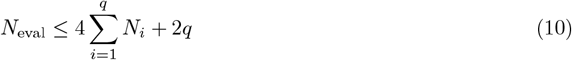

*interval evaluations of* **f**. *Consequently, the worst-case runtime satisfies*

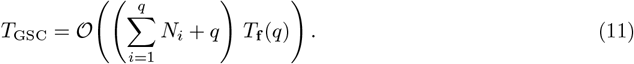

*Proof*. Fix a coordinate *i*. Consider the lower-bound shaving loop for coordinate *i*. Each successful shave removes a slice of width 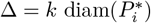 and reduces the width of 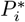 by the factor (1 −*k*). Therefore, the number of successful shaves on that side is at most *N*_*i*_ by Theorem 3. The same bound applies to the upper-bound shaving loop. Hence, across both sides, coordinate *i* admits at most 2*N*_*i*_ successful shaves.

For each shaving attempt, the algorithm evaluates **f** (*P*_test_) once. If the slice is certified infeasible, it additionally evaluates **f** (*P*_prop_) to verify that the proposed remainder remains feasible before committing the contraction. Thus, each successful shave requires at most two interval evaluations. In addition, after the last successful shave on a given side, the loop terminates after one final evaluation of **f** (*P*_test_) that fails to certify removal (or when the slice width falls below *ϵ*). Therefore, each side of coordinate *i* incurs at most 2*N*_*i*_ + 1 evaluations. Summing over both sides gives at most 4*N*_*i*_ + 2 evaluations per coordinate. Summing over *i* = 1, … , *q* gives (10). Multiplying by the per-evaluation cost *T*_**f**_ (*q*) gives (11).

*Remark* 1. The global shaving contractor is certified and contracting: it removes a boundary slice only when interval evaluation certifies that the slice contains no feasible parameter value. As with other interval contractors based on a natural inclusion function, the output is an *outer box enclosure* and may be conservative when interval over-approximation is dominated by dependency effects.

Global shaving is most effective when the steady-state constraints exhibit sufficient *parameter sensitivity* over *X*_*d*_ so that thin boundary slices of *P* can be certified infeasible. This situation commonly arises when (i) the constraints are approximately monotone in one or more parameters over *X*_*d*_, (ii) the parameters are well scaled (so that a fixed shaving ratio *k* corresponds to a meaningful perturbation), and (iii) the inclusion function *F* (*X*_*d*_, ·) remains sufficiently tight on the tested slices. Conversely, if the feasible set is localized away from the boundary of *P*_0_, or if *F* is overly conservative due to repeated variables and strong coupling, the contractor may stall and return a wider enclosure.

Global shaving is *orthogonal* to improvements in range bounding: any tighter inclusion function or evaluation strategy may be substituted for the natural extension used in the case studies. For example, centered/mean-value forms, affine arithmetic, Taylor models, and subexpression-based interval propagation can mitigate dependency and increase the rate at which boundary slices are certified infeasible. The present study uses the natural interval extension to keep guarantees and comparisons transparent; tighter enclosures and propagation-enhanced variants are natural directions for improving contraction quality without introducing subdivision.

## 3 Results

This section demonstrates how the proposed global shaving contractor contracts parameter search domains under design-oriented steady-state specifications. Unless stated otherwise, all inclusion tests use the natural interval extension with outward rounding (Section 2.3.1). All computations were implemented in Julia 1.11.6 using IntervalArithmetic.jl (version 0.20.3); the code is available online [20]. For all examples, the shaving ratio is set to *k* = 10^*−*3^. The tolerance is chosen adaptively as

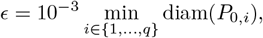

where *P*_0,*i*_ denotes the *i*th coordinate interval of the initial parameter box *P*_0_. For each example, we report the steady-state specification *X*_*d*_, the initial parameter box *P*_0_, the contracted box *P*_pruned_, the contraction ratio vol(*P*_pruned_)*/* vol(*P*_0_), the number of interval evaluations of **f** , and the wall-clock runtime.

### 3.1 Comparison with interval bisection and interval constraint propagation

To quantify the computational overhead of subdivision-based set inversion, we compare the proposed global shaving contractor (Algorithm 1) with two standard interval-analysis baselines: (i) branch-and-bound set inversion based on interval bisection and (ii) interval constraint propagation without subdivision [11, 12, 21].

#### interval bisection (branch-and-bound)

Interval bisection is applied in *parameter space*, starting from *P*_0_ and repeatedly bisecting the widest coordinate of the current box until diam(*P*) ≤ *ϵ*_bisect_. Each candidate box *P* is classified using the same certified exclusion test as in Algorithm 1: if 0 ∉ **f** (*P*) = *F* (*X*_*d*_, *P*), then *P* is discarded; otherwise, *P* is retained as *possibly feasible*. The output is therefore a (generally non-box) certified outer approximation given by a union of retained boxes.

#### Interval constraint propagation

To match the fixed-specification setting of global shaving and interval bisection, *X*_*d*_ is fixed, and interval constraint propagation is applied *only in parameter space*. Specifically, starting from *P*_0_, a forward–backward contractor is iterated on the parameter box using the inclusion map **f** (*P*) = *F* (*X*_*d*_, *P*) and its subexpression structure to contract *P* , without introducing any bisection. As with global shaving, this baseline produces a *certified outer enclosure*: parameters are removed only when interval evaluation certifies infeasibility.

#### Metrics

We report wall-clock runtime, an outer-approximation volume ratio, and the number *N*_eval_ of interval evaluations of the inclusion map **f** (*P*) = *F* (*X*_*d*_, *P*). For methods returning a single box (global shaving and interval constraint propagation), we report the contraction ratio vol(*P*_pruned_)*/* vol(*P*_0_). For interval bisection, we report the normalized retained volume (∑_*j*_ vol(*P*_*j*_) */* vol(*P*_0_). All methods use the same interval extension and outward rounding.

Table 1 shows that global shaving achieves certified contraction without subdivision and requires orders of magnitude fewer inclusion-function evaluations than interval bisection. Across all examples, global shaving reduces *N*_eval_ from millions–billions (interval bisection) to a few thousand, while attaining volume ratios comparable to, or smaller than, the subdivision-based outer approximation. Relative to interval constraint propagation (no bisection), global shaving is typically competitive in runtime and can yield tighter certified enclosures (e.g., Synthetic IFFL and Integral feedback).

**Table 1:**
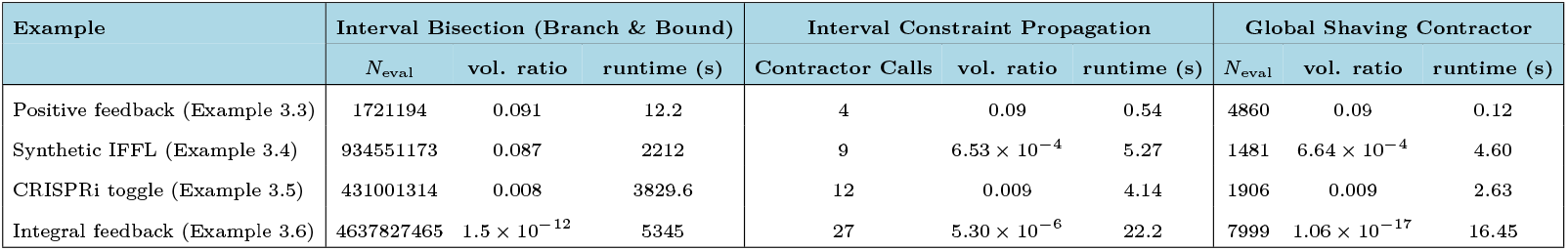
Comparison against interval bisection (branch-and-bound), interval constraint propagation, and global shaving contractor on all examples (global shaving ratio *k* = 0.001). Here, *N*_eval_ denotes the number of interval evaluations of the inclusion map **f** (*P*) = *F* (*X*_*d*_, *P*). The reported vol. ratio is the normalized outer-approximation volume, i.e., vol(*P*_pruned_)*/* vol(*P*_0_) for methods that return a single contracted box and (∑_*j*_ vol(*P*_*j*_) */* vol(*P*_0_) for interval bisection (total retained volume of the union of boxes). The interval-bisection volume ratio is computed at bisection tolerance *ϵ*_bisect_. For Positive feedback, *ϵ*_bisect_ = 0.1 min_*i*_ diam(*P*_0,*i*_). For Synthetic IFFL and Integral feedback, *ϵ*_bisect_ = min_*i*_ diam(*P*_0,*i*_). For CRISPRi toggle, *ϵ*_bisect_ = 0.01 min_*i*_ diam(*P*_0,*i*_).

### 3.2 Effect of inclusion functions

The results above use the natural interval extension. To illustrate the impact of inclusion quality (Section 2.3.1), we additionally run global shaving using a mean-value (centered) enclosure of *g*. Specifically, for each inclusion-function call we evaluate the mean-value form over *Z* = *X*_*d*_ *× P* ,

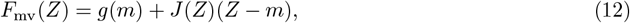

where *m* is the midpoint of *Z* and *J* (*Z*) is an interval extension of the Jacobian of *g* over *Z*. Table 2 reports the resulting contraction and evaluation counts.

**Table 2:** Effect of inclusion quality on global shaving.

| Example | Inclusion $F$ | $N_{\text{eval}}$ | vol. ratio | runtime (s) |
| --- | --- | --- | --- | --- |
| Positive feedback | Natural extension | 4860 | 0.0893 | 1.61 |
|  | Mean-value (centered) | 4742 | 0.0948 | 8.64 |
| Synthetic IFFL | Natural extension | 5618 | 0.0608 | 6.72 |
|  | Mean-value (centered) | 5400 | 0.0678 | 10.97 |

Across both circuits, the mean-value (centered) enclosure yields contraction comparable to the natural extension, but at higher computational cost. For Positive feedback, the mean-value form slightly increases the retained-volume ratio at the expense of runtime. For Synthetic IFFL, a similar trend is observed. Overall, these results illustrate the trade-off between inclusion tightness and per-evaluation overhead: for these examples, the additional derivative computations do not translate into improved end-to-end performance.

We next consider low-dimensional benchmark motifs from prior work on interval-based biomolecular design [21]. The objective is to contract an initial parameter box *P*_0_ while preserving all parameters for which the steady-state constraints admit a solution within the prescribed steady-state specification *X*_*d*_.

### 3.3 Positive feedback loop

A scalar Hill-type positive-feedback model is considered [2, 7].

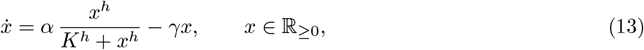

Here *α >* 0 is the maximal production rate, *γ >* 0 is the first-order degradation/dilution rate, *K >* 0 is the half-saturation constant, and *h* is the Hill coefficient. We fix *h* = 10 and define the design vector as *p* = (*α, γ, K*). All inclusion tests are evaluated over the prescribed steady-state specification *X*_*d*_. The steady-state feasibility constraint is

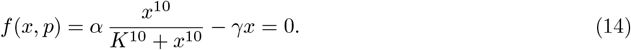

We apply the global shaving contractor to contract the initial parameter search domain using certified infeasibility tests based on interval evaluation. For this model, the steady-state set is disconnected and admits two separated equilibrium intervals. Accordingly, we impose a multistability-oriented specification that requires the existence of one steady state in each of two disjoint desired intervals,

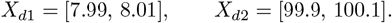

By Theorem 1, we contract the parameter box with respect to each desired interval and intersect the resulting enclosures. Starting from

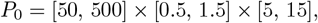

pruning with respect to *X*_*d*1_ yields

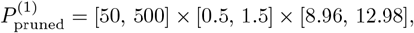

whereas pruning with respect to *X*_*d*2_ yields

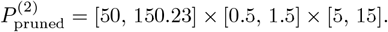

Intersecting these enclosures produces the final contracted box

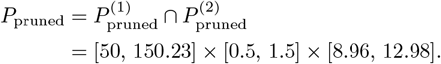

Figure 1 summarizes the initial domain *P*_0_, the intermediate enclosures 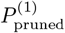 and 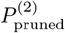 , and their intersection *P*_pruned_.

**Figure 1:**
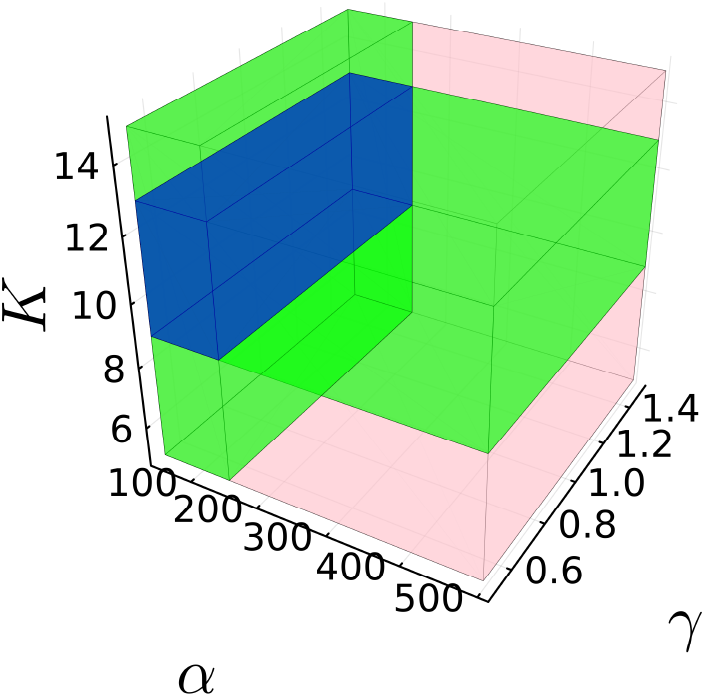
Positive feedback loop design example illustrating interval pruning in parameter space. The pink box is the initial parameter search domain *P*_0_ = [50, 500] × [0.5, 1.5] × [5, 15]. The two green boxes are the pruned enclosures obtained by enforcing the desired steady-state intervals *X*_*d*1_ = [7.99, 8.01] and *X*_*d*2_ = [99.9, 100.1], namely 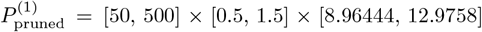 and 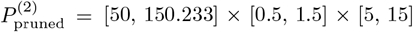. The blue box is their intersection, i.e., the common contracted parameter box returned by the proposed method, *P*_pruned_ = [50, 150.233] *×* [0.5, 1.5] *×* [8.96444, 12.9758].

### 3.4 Synthetic incoherent feedforward loop

A synthetic incoherent feedforward loop, adapted from [15], is described by the nonlinear three-state dynamics

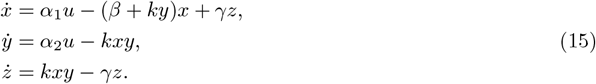

In the design-oriented feasibility setting, the kinetic parameters are treated as decision variables and are chosen such that the system admits a steady state within a prescribed specification *X*_*d*_. We define the design vector as *p* = (*α, β, γ, k*) under the symmetry constraint *α*_1_ = *α*_2_ = *α*, and we apply the global shaving contractor to contract the initial parameter search box. For this example, the desired steady-state interval box is

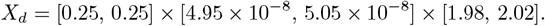

The initial parameter box and the contracted box returned by pruning are

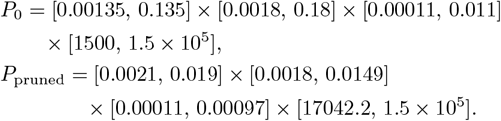

### 3.5 CRISPRi toggle switch model

We consider a CRISPR interference (CRISPRi) toggle switch in which two guide RNAs (sgRNAs), denoted by *s*_1_ and *s*_2_, mutually repress each other through dCas9-mediated transcriptional repression. The objective is to compute certified parameter intervals that guarantee the existence of two distinct steady states within prescribed concentration intervals, corresponding to the two complementary “ON/OFF” configurations of a bistable switch. Models of this form are widely used for the analysis and design of multistable CRISPRi circuit architectures [24].

The sgRNA dynamics are

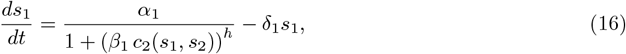

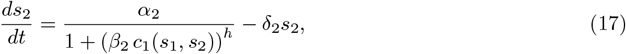

where *s*_1_, *s*_2_ ≥ 0 denote sgRNA concentrations. The production terms model repressed transcription with Hill coefficient *h*, and the linear terms represent first-order degradation/dilution.

Competition for a finite dCas9 pool is represented through the effective complex concentrations

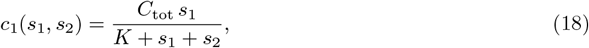

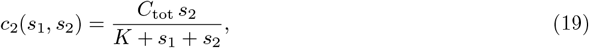

which correspond to a quasi-steady allocation of total dCas9 concentration *C*_tot_ across the two sgRNAs.

Here *α >* 0 is the maximal sgRNA production rate, *δ >* 0 is the degradation/dilution rate, and *β >* 0 quantifies repression strength. The constants *C*_tot_ *>* 0 and *K >* 0 denote the total available dCas9 and an effective dissociation/competition constant, respectively. The nonlinear coupling induced by *c*_1_ and *c*_2_, together with Hill-type repression, can yield bistability for suitable parameter regimes.

#### Design specification and contracted parameter box

To encode bistability, we impose a simultaneous steady-state specification consisting of two disjoint desired steady-state boxes, *X*_*ss*_ = *X*_*d*1_ ∪ *X*_*d*2_, where

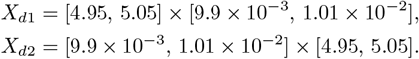

This specification corresponds to the two complementary expression patterns: (*s*_1_, *s*_2_) high/low and low/high.

We fix *h* = 5 and impose a symmetric parameterization,

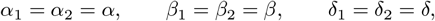

with a common dCas9 competition constant *K*. Accordingly, the design vector is *p* = (*α, β, δ, K*). In all computations, we fix *C*_tot_ = 5 and evaluate inclusion tests over the prescribed specification *X*_*d*_.

Starting from the initial search box (ordered as *α × β × δ × K*)

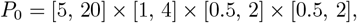

the contractor returns the pruned boxes

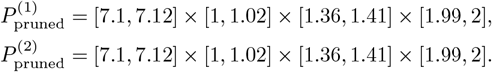

In this case, 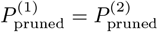 , and hence the final contracted box is

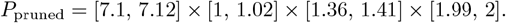

### 3.6 Synthetic biomolecular integral controller

Integral control has been shown to underlie homeostasis and adaptation in a variety of biomolecular contexts [2, 7]. The synthetic integral biomolecular controller model from [1] is considered, yielding a nine-dimensional nonlinear system and a sixteen-parameter design vector. The dynamics are

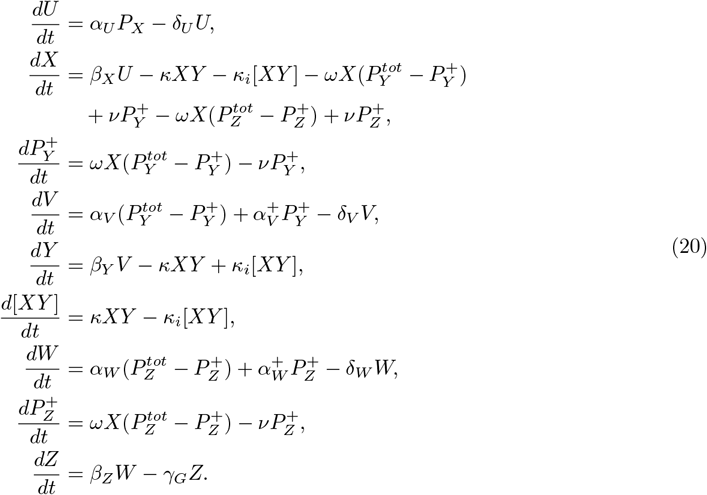

Here *x* = [*U, X*, 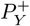, *V, Y*, [*XY*], *W*, 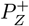, *Z*] *∈* ℝ^9^ and *p* = [*α*_*U*_ , *δ*_*U*_ , *β*_*X*_ , *κ, κ*_*i*_, *ω, ν, α*_*V*_ , 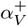, *δ*_*V*_ , *β*_*Y*_ , *α*_*W*_ , 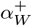 , *δ*_*W*_ , *β*_*Z*_, *γ*_*G*_] *∈* ℝ^16^ denote the state and parameter vectors. The desired steady-state interval is specified as

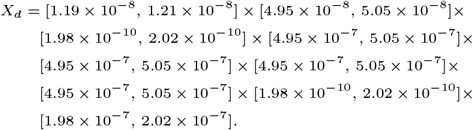

The constants are fixed to *P*_*X*_ = 0.5 nM, 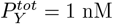, and 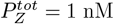. For the initial parameter search box *P*_0_, the proposed pruning procedure contracts it to *P*_pruned_. Table 3 reports the parameter-wise intervals.

**Table 3:**
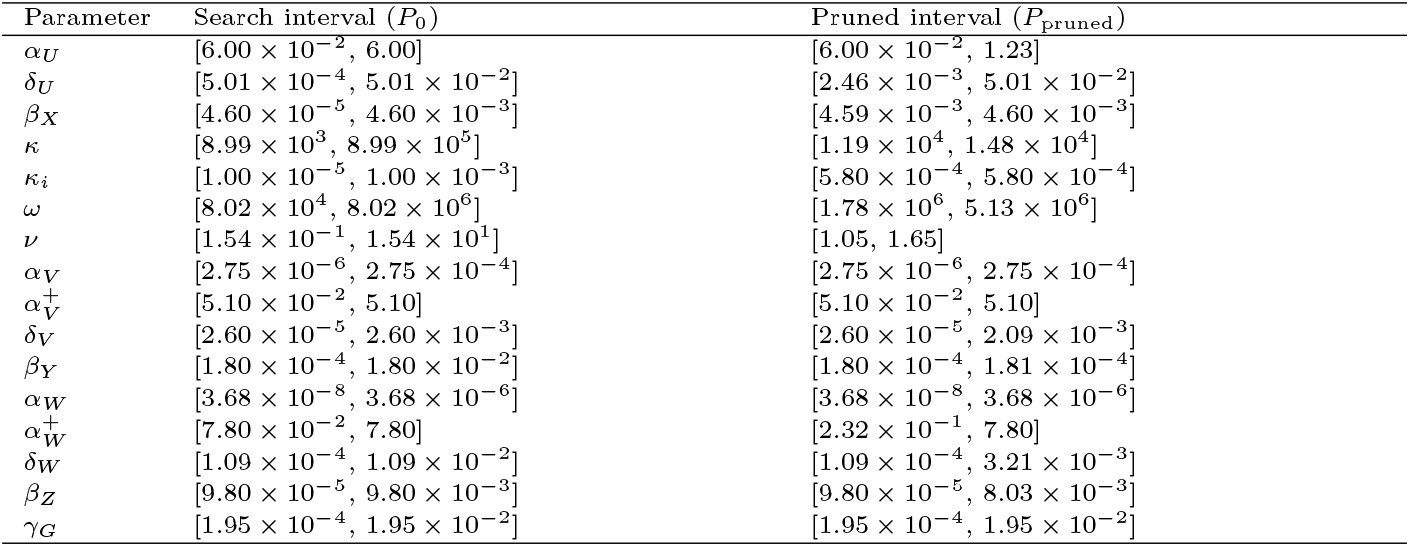
Synthetic biomolecular integral controller: initial parameter search intervals (*P*_0_) and the contracted intervals (*P*_pruned_) returned by pruning.

## 4 Discussion

The case studies indicate that global shaving can compute certified parameter-interval enclosures for steady-state specifications and can substantially contract initial parameter boxes. For the low-dimensional motifs, pruning removes large boundary regions of *P*_0_ and yields markedly tighter enclosures. For the CRISPRi toggle switch design, the prune-and-intersect workflow enforces two disjoint steady-state specifications corresponding to the complementary expression patterns.

The integral-feedback example illustrates the behavior in a higher-dimensional setting. Several parameters contract to narrow intervals (e.g., *β*_*X*_ and *κ*_*i*_), whereas others remain only weakly constrained by the selected steady-state specification. This heterogeneity is consistent with strong coupling and interval dependency: exclusion tests may be decisive along some parameter directions while remaining inconclusive along others, and the overall computational cost can be sensitive to the tuning parameters.

From a practical standpoint, the contractor requires only an inclusion function for the steady-state constraints and returns a guaranteed outer enclosure of the feasible parameter set. Because *X*_*d*_ is fixed, the computation focuses on contracting *P*_0_ rather than exploring the state space. Moreover, the procedure is compositional and can be applied repeatedly to encode multiple steady-state requirements.

As a boundary-based contraction method, global shaving may provide limited contraction when the feasible set is strongly disconnected or localized away from the boundary of *P*_0_. In strongly coupled and higher-dimensional models, interval dependency and wrapping may prevent exclusion on slices that are mostly infeasible, thereby increasing the number of evaluations required to reach a prescribed tolerance.

The same inclusion-based framework also applies to the complementary *analysis* problem: for a fixed parameter box *P*_0_, one can compute guaranteed steady-state bounds in state space by contracting an initial state box (or a collection of boxes) using inclusion tests on *F* (*X, P*_0_). In addition, integrating global shaving with limited subdivision (branch-and-prune) can mitigate dependency by shrinking boxes prior to evaluation, potentially improving pruning at the expense of additional evaluations. Future work includes benchmarking against interval-Newton and advanced propagation-based contractors, characterizing sensitivity to (*k, ϵ*), and extending the specification to include local stability, robustness margins, and transient performance requirements.

## 5 Conclusions

We presented a global shaving contractor for certified steady-state parameter-interval design in biomolecular models with bounded uncertainty. Using interval inclusion tests, the method removes boundary regions of the parameter domain that cannot satisfy prescribed steady-state constraints while maintaining a guaranteed outer enclosure of the feasible parameter set. We also provided certification and complexity results that characterize computational cost in terms of the number of inclusion-function evaluations. In the case studies, global shaving reduced the number of inclusion-function evaluations from 1.7 ×10^6^– 4.6× 10^9^ (interval bisection) to 1.5 ×10^3^–8.0 ×10^3^, while achieving contracted-volume ratios as low as 1.06 ×10^*−*17^ in the sixteen-parameter example. These guarantees support certified parameter screening and uncertainty-aware steady-state design workflows in which infeasible regions of *P*_0_ are excluded without subdivision while preserving a rigorous outer enclosure of all feasible designs.

